# Environmental exosomes: Evidence of extracellular RNA release by aquatic organisms

**DOI:** 10.1101/2025.11.04.686478

**Authors:** Ryo Yonezawa, Lingxin Meng, Naoki Hashimoto, Ibuki Igarashi, Satoshi Kimura, Nina Yasuda, Susumu Mitsuyama, Takanori Kobayashi, Kazutoshi Yoshitake, Shigeharu Kinoshita, Nahoko Bailey-Kobayashi, Kaoru Maeyama, Kiyohito Nagai, Shugo Watabe, Tetsuhiko Yoshida, Shuichi Asakawa

**Affiliations:** Laboratory of Aquatic Molecular Biology and Biotechnology, Department of Aquatic Bioscience, Graduate School of Agricultural and Life Science, The University of Tokyo, Bunkyo, Tokyo 113-8657, Japan; Signal Peptidome Research Laboratory, Department of Aquatic Bioscience, Graduate School of Agricultural and Life Sciences, The University of Tokyo, Bunkyo, Tokyo 113-8657, Japan; Laboratory of Aquatic Conservation, Department of Ecosystem Studies, Graduate School of Agricultural and Life Science, The University of Tokyo, Bunkyo, Tokyo 113-8657, Japan; Technology Advancement Center, Graduate School of Agricultural and Life Sciences, The University of Tokyo, Bunkyo-ku, Tokyo 113-8657, Japan; College of Bioresource Sciences, Nihon University, Kanagawa, 252-0880, Japan; School of Marine Biosciences, Kitasato University, Minami-ku, Sagamihara, Kanagawa 252-0313, Japan; Institute for Advanced Sciences, TOAGOSEI CO., LTD., Tsukuba, Ibaraki 300-2611, Japan; Mikimoto Pearl Research Institute, K.MIKIMOTO & CO., LTD., Hazako 923-74, Hamajima, Shima, Mie 517-0403, Japan; Mikimoto Pharmaceutical CO., LTD., Kurose 1425, Ise, Mie 516-8581, Japan

## Abstract

Aquatic organisms continuously interact with surrounding water, yet whether they release extracellular vesicles remains unknown. We hypothesized that pearl oysters (*Pinctada fucata*) release exosomes/small extracellular vesicles (sEVs) into the aquatic environment. To this end, we collected exosomes/sEV-sized components by ultrafiltration from tank water and open-sea culture areas. Microscopy revealed abundant vesicles consistent with exosome/sEV size, and RNA sequencing identified oyster-specific piRNAs that matched sequences previously detected in hemolymph exosomes. These findings demonstrated that pearl oysters actively released exosomes containing species-specific nucleic acids into surrounding water. We propose referring to these vesicles as environmental exosomes/environmental sEVs (eExosomes/esEVs). This finding suggests that aquatic exosomes serve as carriers of RNA and may contribute to inter-organismal communication networks. Beyond their functional role, eExosomes/esEVs also hold promise as novel targets for environmental DNA/RNA (eDNA/eRNA) analysis, offering new opportunities for ecological monitoring and biodiversity research.

## Introduction

Aquatic organisms continuously exchange biological components with their surrounding waters. These exchanges can profoundly influence ecological interactions. We hypothesized that, in addition to soluble biomolecules, exosomes and small extracellular vesicles (sEVs) may also be secreted into aquatic environments. These vesicles could act as stable carriers of biological information not only within organisms (as they do in terrestrial species, transferring information among tissues and organs), but also between organisms. Exosomes and sEVs are lipid bilayer vesicles typically 50–200 in diameter (*1*–*6*), known to encapsulate nucleic acids, proteins, and other bioactive molecules (*2, 7*–*9*) and mediate a wide range of intercellular communication processes (*2, 3, 7, 10*). In terrestrial and biomedical research, extensive studies have revealed their critical roles in physiological regulation, disease mechanisms, and cell-to-cell signaling (*11*–*15*). However, investigations into the roles of exosomes in aquatic organisms remain limited. Most prior work has focused on intracellular or organismal functions, while their potential release into the external aquatic environment and possible contributions to inter-organismal communication remain essentially unexplored.

Extracellular vesicles, such as exosomes, possess a lipid bilayer that protects their enclosed nucleic acids and proteins from enzymatic degradation (*16*), potentially enabling their persistence in aquatic environments. If released into surrounding waters, these vesicles could serve as vehicles for the transfer of functional molecules between individuals, thus contributing to ecological communication networks. Our previous studies on the Akoya pearl oyster (*Pinctada fucata*) have demonstrated that hemolymph-derived exosomes are enriched in small RNAs, particularly PIWI-interacting RNAs (piRNAs) (*17, 18*), which are expressed in various somatic tissues of invertebrates (*19, 20*). In model organisms, such as humans and *Drosophila*, piRNAs are predominantly expressed in reproductive tissues, where their primary function is the suppression of transposable elements (*21, 22*). In contrast, invertebrate piRNAs appear to serve broader functions, potentially resembling those of microRNAs (miRNAs) (*19, 23*). Empirical evidence supports this possibility; for instance, plant-derived microRNAs have been shown to regulate insect gene expression (*24*), and algae-derived exosomes and miRNAs have been detected in *P. fucata* (*25*). These findings suggest that small RNA-mediated communication, including regulatory functions, may occur in aquatic environments, with exosomes acting as protective and transmissive carriers for these molecules.

In recent years, environmental nucleic acids (eNAs) have emerged as powerful tools for assessing biodiversity and conducting ecological monitoring (*26*–*29*). However, the inherent instability of RNA under natural conditions limits its use in field applications (*30, 31*). Addressing this challenge requires identifying alternative extracellular carriers that can stabilize RNA under environmental conditions.

Therefore, in this study, we aimed to provide the first direct evidence that Akoya pearl oysters release exosomes and sEVs into surrounding seawater by combining microscopy-based characterization with small RNA sequencing.

## Results

### Identification of exosomes/sEVs from environmental seawater

Exosomes/sEV fraction were isolated from the high-molecular-weight component of filtered environmental seawater using ultrafiltration. One sample from the rearing tank and one each from the morning (AM) and afternoon (PM) aquaculture areas were examined via microscopy. Vesicle size and concentration were measured using interferometric light microscopy (Videodrop; Myriade, Paris, France). The median diameters of the AM and PM samples were 176 nm and 174 nm, respectively (Fig. 1). The concentrations were approximately 5.9 × 10^8^ and 8.0 × 10^8^ vesicles/mL, respectively. These values were similar to those reported for exosomes derived from Akoya pearl oyster tissue and hemolymph in previous studies by Meng et al. (*32*) and Huang et al. (*17*), as well as to preliminary data from rearing tank water (data not shown). When extrapolated to 1 L of environmental seawater, the estimated vesicle concentration ranged from approximately 1.0 × 10^10^ to 10^11^ vesicles/L. Transmission electron microscopy (TEM) revealed numerous vesicles with diameters primarily in the 50–200 nm range. In addition, vesicles smaller than 50 nm and filamentous structures were observed that were not detectable via interferometric light microscopy (Fig. 2A and B).

**Fig. 1.**
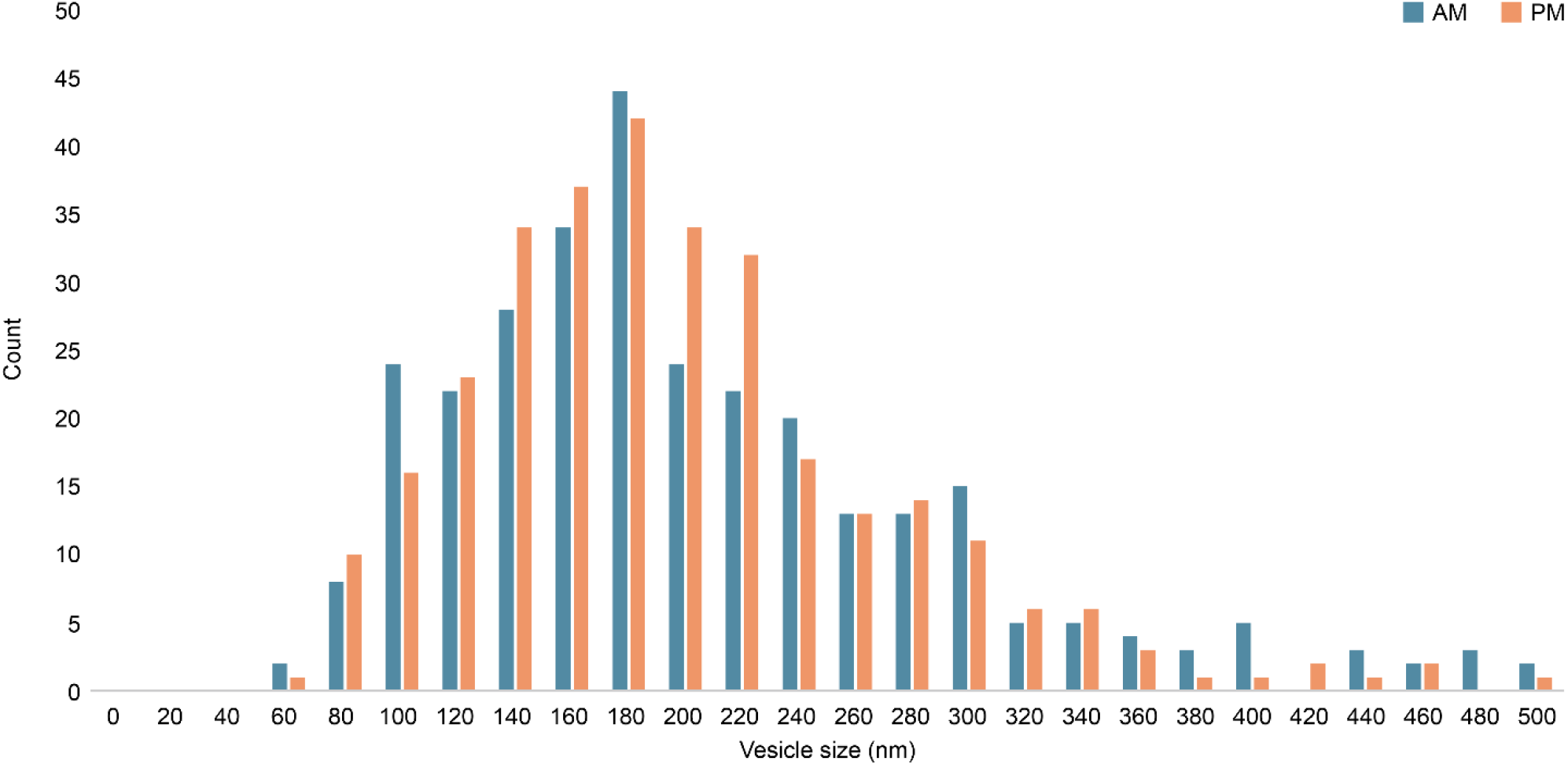
Exosome/sEV fraction identification using interferometric light microscopy of Videodrop. Vesicle size distribution of exosomes/sEVs isolated from open-water samples collected in the morning (AM) and evening (PM).

**Fig. 2.**
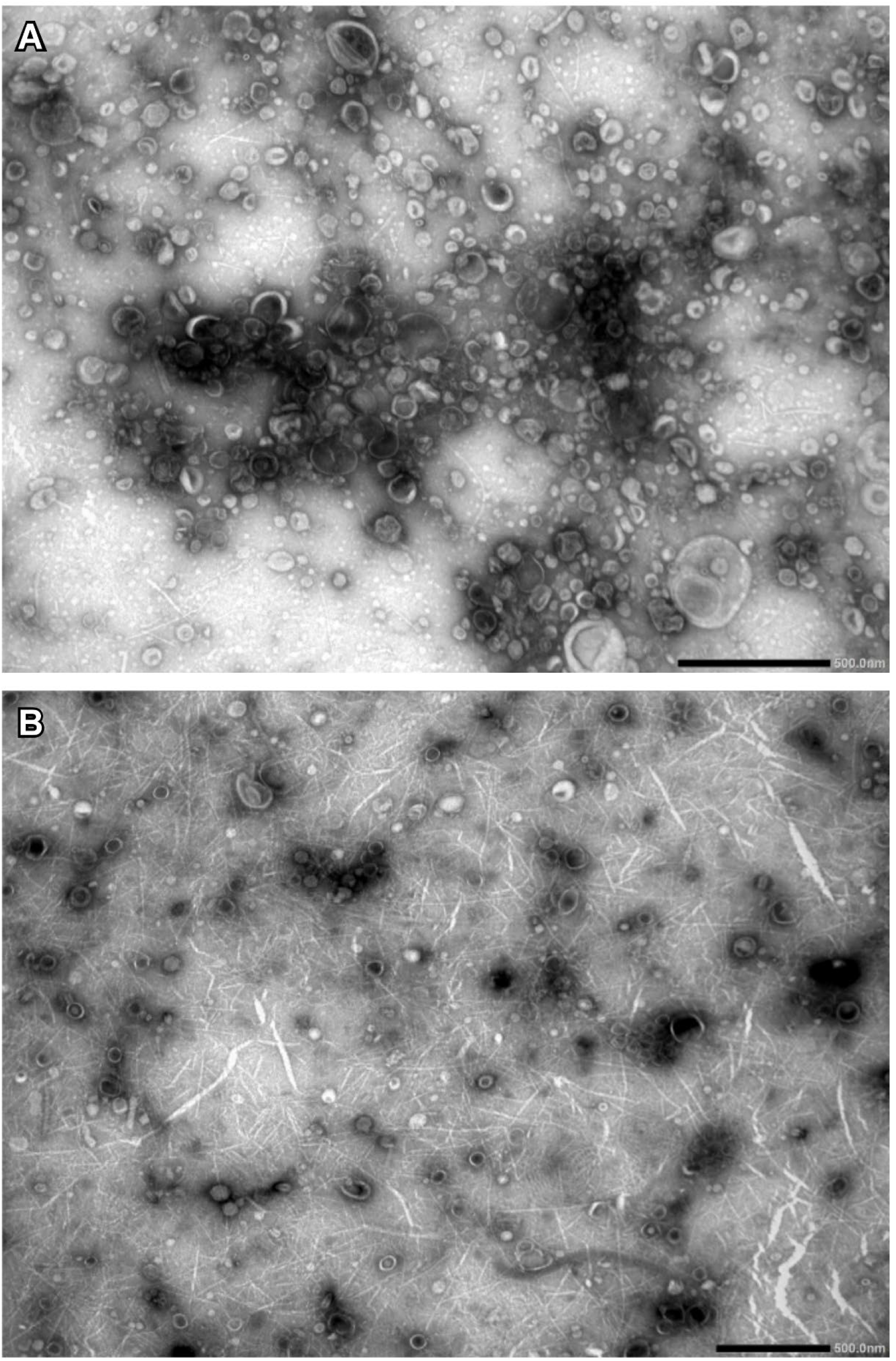
Transmission electron microscopy (TEM) images of exosome/sEV fractions. (A) Exosomes/sEVs isolated from tank water. (B) Exosomes/sEVs isolated from morning (AM) open water sample. Spherical vesicles are indicated; vesicles smaller than 50 nm and filamentous structures may correspond to ribosomes and nucleic acids, respectively. Scale bars: 500 nm.

Small RNA was extracted from aquaculture water samples collected at the same site in the morning (AM, n=2) and in the afternoon (PM, n=2), and once from the rearing tank sample using the exoRNeasy kit (Qiagen, Hilden, Germany). The resulting exosome/sEV preparations were then quantified using the Qubit microRNA assay. Adequate RNA yields were obtained, with concentrations of 9.0 ng/μL for the rearing tank sample and 7.2 ng/μL (AM1), 15.2 ng/μL (AM2), 11.1 ng/μL (PM1), and 20.6 ng/μL (PM2) for the aquaculture samples. Electrophoretic analysis using an Agilent Bioanalyzer showed that the majority of RNA fragments were shorter than 40 nucleotides. In most samples, more than 70% of sequences fell within the characteristic size range for miRNAs.

### Small RNA Profiling of Exosomes/sEVs

Duplicate samples were collected from the aquaculture area in the morning and afternoon, and one sample was collected from the rearing tank. These samples were subjected to small RNA sequencing using the DNBSEQ-G400 platform (BGI, HongKong, China) with 100 bp paired-end reads. The raw sequencing output for the rearing tank and aquaculture samples (AM1, AM2, PM1, and PM2) was 40 million, 30 million, 8 million, 13 million, and 9 million reads (Mreads), respectively. After adapter trimming and quality filtering, reads were restricted to a length of 18–40 nucleotides (nt). The remaining read counts were: rearing tank, 15 Mreads; AM1, 13 Mreads; AM2, 4 Mreads; PM1, 6 Mreads; PM2, 5 Mreads. Mapping these reads to the *P. fucata* reference genome resulted in alignment rates of 4.6% for the tank water sample and 2.8 ± 0.2% for aquaculture area samples. Even though nearly two days elapsed between seawater collection and vesicle isolation, a considerable proportion of reads still mapped to the P. fucata reference genome.

Reads were annotated by mapping to the Rfam and miRbase databases. Sequences of 20– 24 nt were classified as canonical miRNAs, while sequences of 25–32 nt that did not map to known small RNA categories were designated as estimated piRNAs. Annotated reads were further categorized into piRNA, miRNA, rRNA, tRNA, snRNA, other ncRNAs, and unknown RNA types. Corresponding read counts for each category are presented in Table 1. Among these, rRNA was the most abundant RNA type across all samples (Fig. 3). In contrast, miRNAs accounted for less than 0.1% of mapped reads. Notably, despite being extracted from environmental seawater, piRNAs were detected in higher-than-expected proportions, 2.5% in the rearing tank sample and 0.1 ± 0.06% in aquaculture samples.

**Table 1.** RNA type distribution based on small RNA-seq mapping and annotation.

| **RNA/Reads** | **Tank** | **AM1** | **AM2** | **PM1** | **PM2** |
| --- | --- | --- | --- | --- | --- |
| **piRNA** | **19,248** | **372** | **81** | **246** | **145** |
| **miRNA** | **27** | **30** | **9** | **22** | **7** |
| **rRNA** | **373,789** | **361,592** | **103,368** | **171,123** | **107,696** |
| **tRNA** | **45,384** | **4,174** | **1,606** | **4,321** | **2,328** |
| **snRNA** | **1,239** | **1,638** | **589** | **717** | **1,347** |
| **other_ncRNAs** | **2,293** | **830** | **173** | **342** | **1,088** |
| **unknown** | **338,564** | **8,702** | **3,025** | **5,376** | **3,306** |

**Fig. 3.**
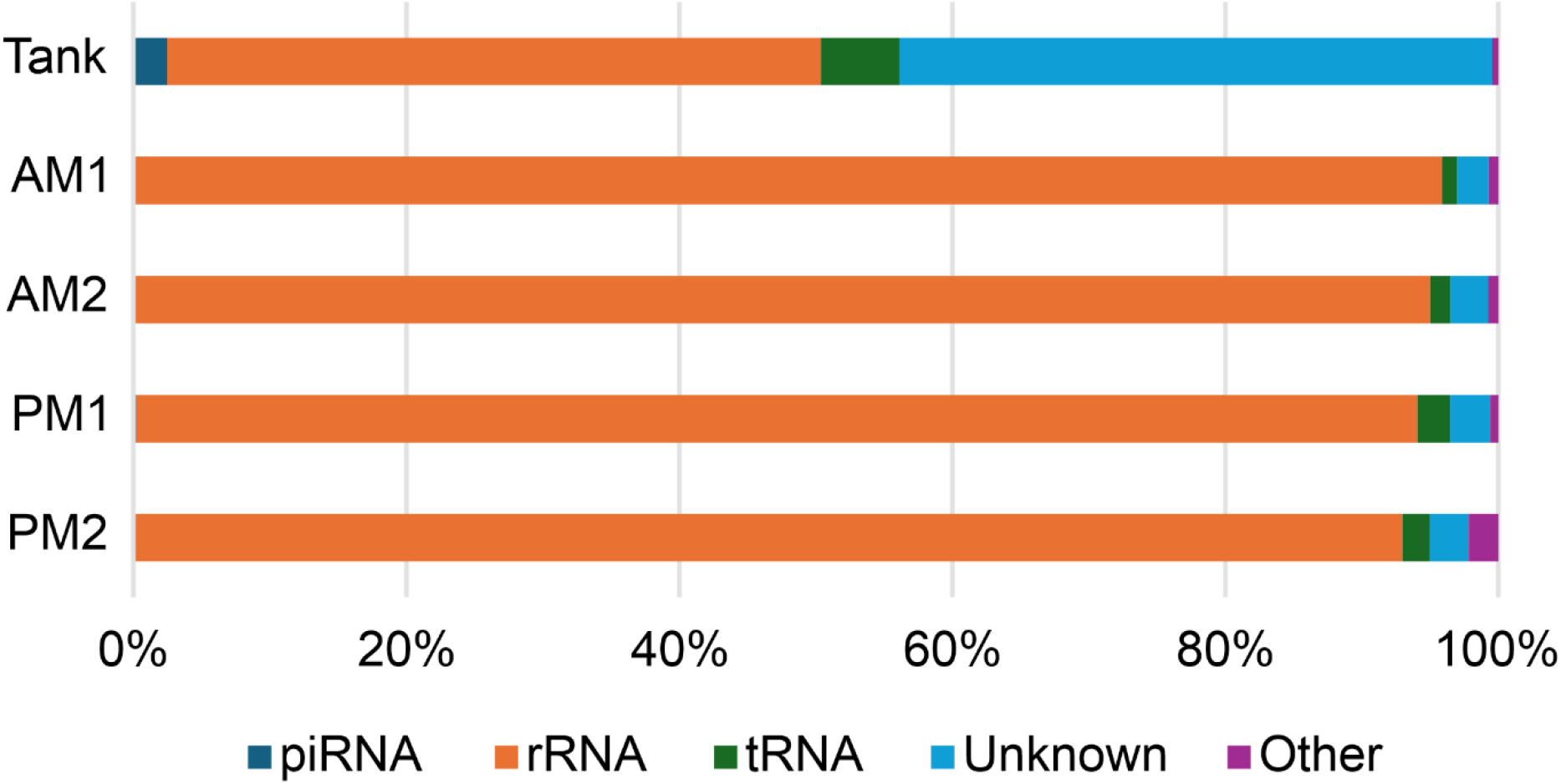
Classifications of mapped small RNA reads from rearing tank and aquaculture water samples. piRNA, piwi-interactive RNA; rRNA, ribosomal RNA; tRNA, transfer RNA; unknown, non-annotated sequence; Other, other small RNAs.

The sequence most frequently identified among the estimated piRNAs was observed 251 times in the rearing tank samples and 33 times in aquaculture samples (Supplementary data 1). The 1U bias (*33*–*35*), calculated based on the number of estimated piRNA sequence types, was 31% in the rearing tank and 22.4 ± 2.1% in the aquaculture area. The corresponding frequency rates were 19.1% and 15.4 ± 5.8%, respectively (Table 2).

**Table 2.** 1U bias based on the number of sequence types and observed frequencies.

| Sample | Types of sequence |  |  | Frequencies of sequence |  |  |
| --- | --- | --- | --- | --- | --- | --- |
|  | U | A, G, C | 1U bias (%) | U | A, G, C | 1U bias (%) |
| Tank | 863 | 2,067 | 31.3 | 3,679 | 15,569 | 19.1 |
| AM1 | 71 | 230 | 23.6 | 74 | 298 | 19.9 |
| AM2 | 16 | 51 | 23.9 | 16 | 65 | 19.8 |
| PM1 | 27 | 112 | 19.4 | 35 | 211 | 14.2 |
| PM2 | 8 | 27 | 22.9 | 11 | 134 | 7.6 |

Sequence-type-level analysis indicated that a 1U bias was present only in the rearing tank samples. Comparison of the estimated piRNAs with exosome-derived piRNAs reported by Huang et al. (*17*), using BLASTn, revealed that 116 sequence types from the rearing tank and 6 types from the aquaculture area matched exactly with previously reported sequences. Several piRNAs frequently detected in earlier studies were also found in this dataset. Among these, the most abundant piRNA was observed 198 times in the rearing tank sample and 13 times in the aquaculture water sample (Tables 3 and 4; Supplementary data 2).

**Table 3.**
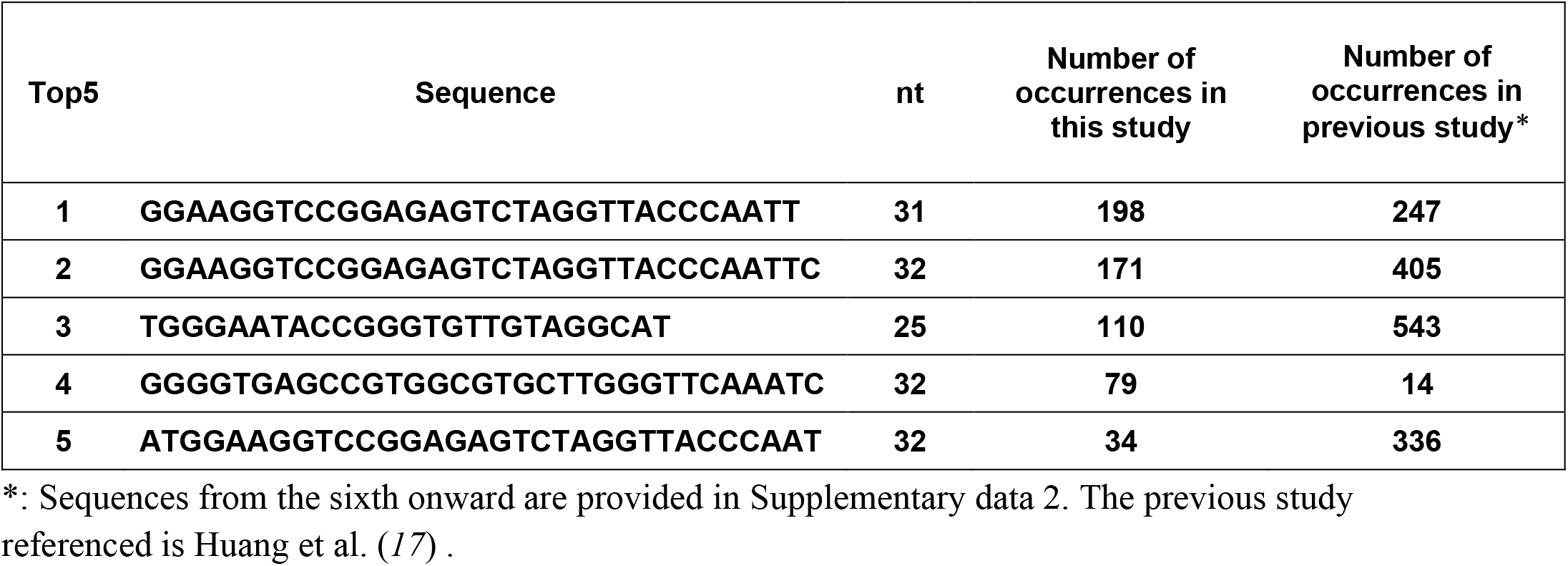
Frequently detected piRNA sequences in the rearing tank samples.

**Table 4.**
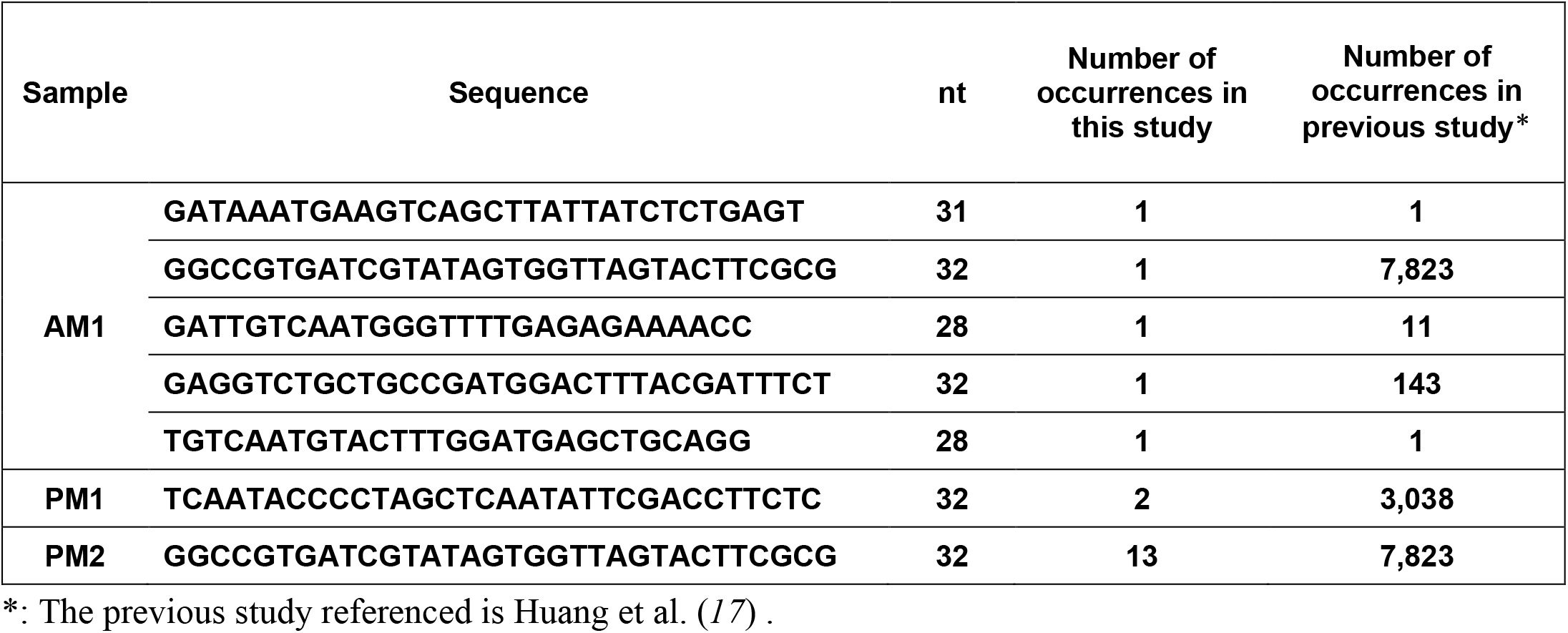
Occurrence counts of piRNA sequences matching known sequences in aquaculture water samples.

### mRNA detection from small RNA-seq data

mRNA analysis was performed using paired-end reads from the rearing tank and AM1 aquaculture samples, both of which yielded sufficient sequencing depth. After filtering, reads were aligned to the *P. fucata* reference genome. The number of paired-end reads mapped was 56 Kreads (Tank) and 120 Kreads (AM1). Transcript assembly was performed using the Hisat2– StringTie–Trinotate pipeline. This analysis yielded 164 transcripts from the rearing tank (125 of which were annotated) and 49 transcripts from AM1 (10 annotated), including transcripts lacking gene annotations (Supplemental data 3). Among the detected transcripts were lysosome-associated membrane glycoprotein 1 (LAMP1) (*36*), moesin (MOES) (*37*), BRO1 domain-containing protein BROX (BROX) (*38*), and filamin-A (FLNA) (*39*), all of which have previously been reported in exosomes/EVs. Additionally, several genes associated with key cellular functions were detected, including those involved in signal transduction (Signal transducer and activator of transcription 5A; STAT5A), reproduction (Spermatogenesis-associated serine-rich protein 2;SPAS2, Zonadhesin;ZAN), and transport (ATP-binding cassette sub-family A member 2; ABCA2, Sodium- and chloride-dependent glycine transporter 2; SLC6A5).

## Discussion

Exosomes are small extracellular vesicles secreted by eukaryotic cells that contain nucleic acids, proteins, lipids, and other bioactive molecules (*2, 7, 8*). They have been extensively characterized in bodily fluids and tissues, where they participate in intercellular communication (*40, 41*). However, nearly all prior studies have focused on intracellular or intercellular functions; their release into and potential interaction with environmental seawater remain largely unexplored. Our findings provide the first direct evidence that aquatic organisms, specifically the *P. fucata*, release exosomes into their surrounding aquatic environment. This observation reveals a previously unrecognized biological phenomenon. We propose that these extra-individual exosomes/sEVs be termed “environmental Exosomes/environmental sEVs (eExosomes/esEVs)”.

By adapting an established exosome/sEV isolation protocol originally developed for cell culture supernatants and liquid biopsy samples (*42*), we successfully obtained vesicle fractions from environmental seawater. These fractions exhibited size distributions consistent with those of tissue- and hemolymph-derived exosomes (26, 35), as determined by interferometric light microscopy (Videodrop). Transmission electron microscopy further confirmed the presence of abundant vesicles sized 50–200 nm, which aligns with the known size range of exosomes (*17*). These results demonstrate that vesicles morphologically and physically consistent with standard exosomes are present in environmental seawater and validate the utility of this modified method for environmental exosome isolation.

Small RNA-seq of these vesicle fractions revealed that despite a delay of nearly 2 days between sampling and isolation, a considerable proportion of reads could still be mapped to the *P. fucata* reference genome. This mapping rate was comparable to or exceeded that reported in conventional eRNA analyses, where RNA degradation typically occurs within 24 h (*27, 43*). These findings strongly suggest that the lipid bilayer of exosomes confers protection against enzymatic degradation, microbial digestion, and other environmental stressors, thereby preserving the integrity of encapsulated nucleic acids (*16, 44*). Most of the identified sequences corresponded to rRNA, which may partially reflect the co-isolation of free nucleic acids (*45*) and ribosomes (*46*), as supported by the presence of filamentous structures and vesicles smaller than 50 nm observed via TEM (Fig. 2). As shown in Tables 2 and 3, we also detected piRNAs with sequences identical to those previously identified in hemolymph-derived exosomes (*17*), providing strong evidence that species-specific small RNAs are released into the environment via exosomes. piRNAs are generally characterized by strong species or lineage specificity and are classically associated with transposon silencing in reproductive tissues or organs of model organisms, such as *Drosophila* and humans (*47, 48*). However, in aquatic invertebrates, piRNAs are distributed across various tissues and organs. In the Akoya pearl oyster, piRNAs are not only abundant in reproductive tissues but also enriched in hemolymph-derived exosomes (*19, 20, 23*). Recent studies have implicated piRNAs in additional biological processes, including developmental regulation, immune defense, and gene expression control (*49*–*52*). mRNA analysis using small RNA-seq data identified transcripts for LAMP1 (*36*), MOES (*37*), BROX (*38*), and FLNA (*39*), all of which have previously been shown to be associated with exosomes/sEVs. Among these, LAMP1 is considered an exosome-specific surface protein, FLNA is frequently detected in exosomes, and MOES is regarded as a universal pan-exosome marker (*36, 53, 54*). These findings provide indirect support for the interpretation that the vesicles detected in the environmental seawater samples correspond to exosomes/sEVs. Although their precise association with exosomes remains unclear, the presence of transcripts related to reproduction, transport, and signaling may suggest roles in inter-individual communication.

Collectively, our results support the hypothesis that exosome-mediated release disseminates functional small RNAs and mRNAs beyond individual organisms, potentially influencing processes at the population level. Exosome-encapsulated piRNAs, mRNAs, and other bioactive molecules may participate in inter-individual signaling. For example, they may facilitate immune-related communication by conveying information about viral or bacterial infections to conspecifics, thereby promoting population-level defense mechanisms. Similarly, given the established roles of exosomes in reproduction in other animals (*55*–*57*) and the collective spawning behaviors observed in many marine organisms, such as pufferfish and corals (*58, 59*), exosome release may contribute to the coordination of reproductive events. Although the specific biological functions of eExosomes/esEVs remain to be determined, our quantification of more than 10 billion vesicles per liter of seawater highlights their potential ecological relevance. Given that pheromones and other chemical signals exert biological effects at far lower concentrations (*60*), exosomes may represent a novel and previously unrecognized mechanism for inter-individual or organismal communication in aquatic ecosystems.

Despite these promising findings, our study has several limitations. First, the exosome/sEV fractions isolated from environmental seawater may include co-isolated particles such as ribosome and/or free nucleic acids, which could complicate interpretation of the sequencing data. Second, although piRNAs and mRNAs detected in these vesicles suggest potential biological roles, functional validation of their activity in recipient organisms was beyond the scope of this study. Third, our analyses were limited to a single species and sampling period, and additional studies across multiple organisms and environmental contexts will be required to assess the generality of our observations. Addressing these limitations will be essential to fully elucidate the ecological and physiological significance of eExosomes/esEVs.

In conclusion, this study demonstrates for the first time that aquatic organisms release exosomes containing species-specific nucleic acids into their surrounding waters. By stabilizing and transmitting functional molecules, eExosomes/esEVs may represent a previously unrecognized mode of inter-organismal communication and contribute to the formation of complex environmental information networks. Notably, unlike eRNA, which rapidly degrades under natural conditions, eExosomes/esEVs provide a stable source of molecular information, suggesting that they may complement existing eDNA and eRNA approaches in ecological monitoring. Future research should aim to elucidate the mechanisms, dynamics, and ecological significance of eExosomes/esEVs across diverse aquatic organisms.

## Materials and Methods

### Biological materials

In September 2023, a tank containing Akoya pearl oysters (summer-harvested seed bivalves, 300 individuals in a 30-L tank with 20 L of filtered seawater) was maintained for 5 days under static water conditions without feeding. As the initial findings were derived from closed-environment tank water, we extended our investigation to open-water conditions conducted concurrently. As the initial findings were derived from closed-environment tank water, we extended our investigation to open-water conditions conducted concurrently. In September 2024, 1 L of water was collected twice (total volume: 2 L), once in the morning and once in the evening, from the Akoya pearl oyster aquaculture area (34°17’41”N 136°48’09”E) in Ago Bay, Mie Prefecture, Japan (Fig. 4). After collection, the water samples were stored at 4 °C for 1 day and transported to the University of Tokyo, Tokyo, Japan, via refrigerated delivery at 4 °C. Samples were processed immediately upon arrival. This research was approved by the Animal Experiment Ethics Committee of the Graduate School of Agricultural and Life Science, The University of Tokyo (Accession No. P21-103).

**Fig. 4.**
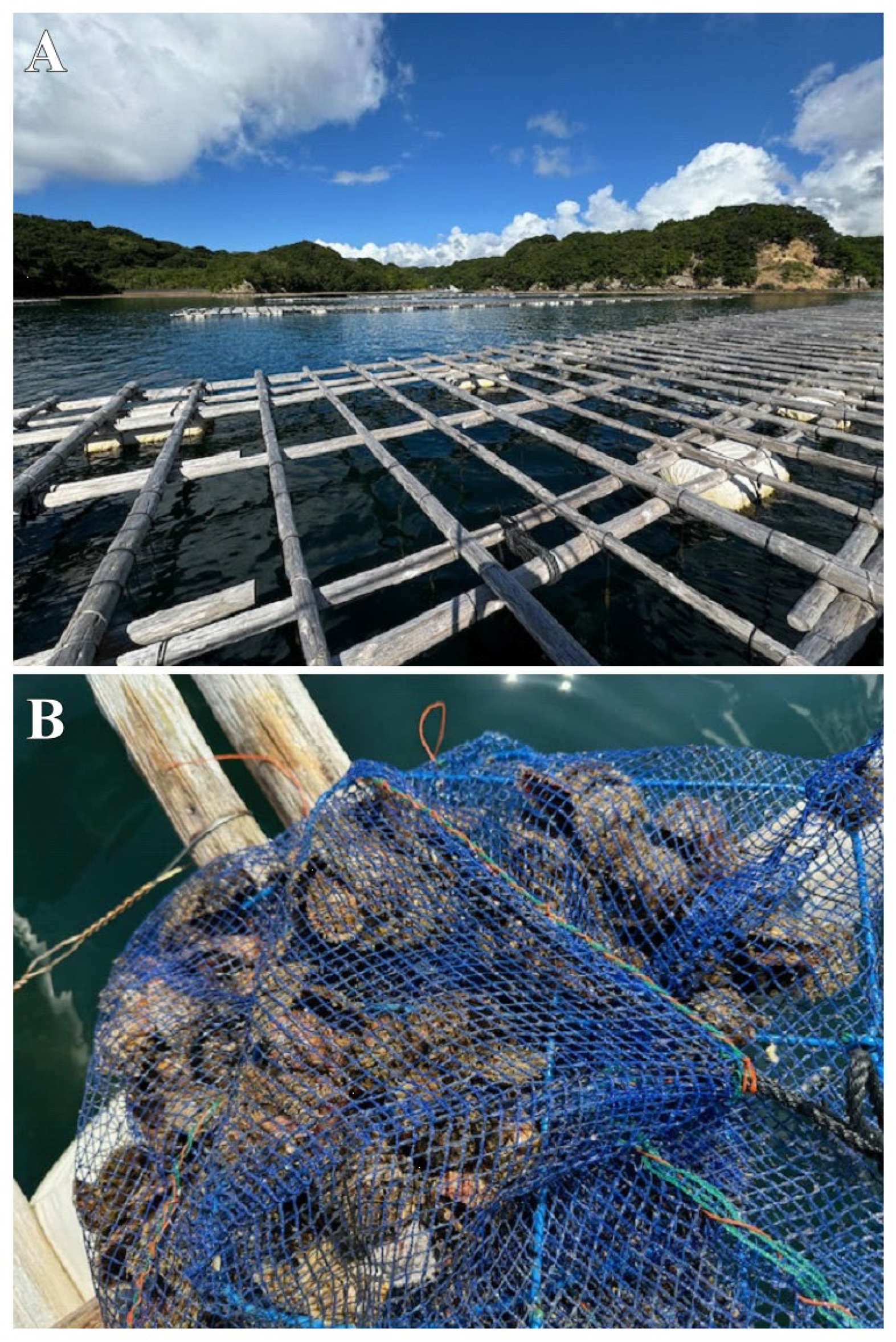
Sampling sites and collection procedures in the aquaculture area. (A): The raft used for farming Akoya pearl oysters in Ago Bay, Shima, Mie prefecture. (B): Suspended net cage for Akoya pearl oyster marine farming.

### Environmental exosome isolation and identification

As shown in Fig. 5, exosomes/sEVs were recovered by modifying an exosome purification protocol originally designed for culture supernatants (*42*). To isolate vesicles within the exosome/sEV size range (50–200 nm), seawater was first filtered through a 20-μm mesh to remove large particulates. The filter was then passed through a 0.22-µm Stericup-GP filter (Merck Millipore, MA, USA) to eliminate residual smaller particulates. The final filtrate, containing vesicles smaller than 220 nm, was subjected to ultrafiltration.

**Fig. 5.**
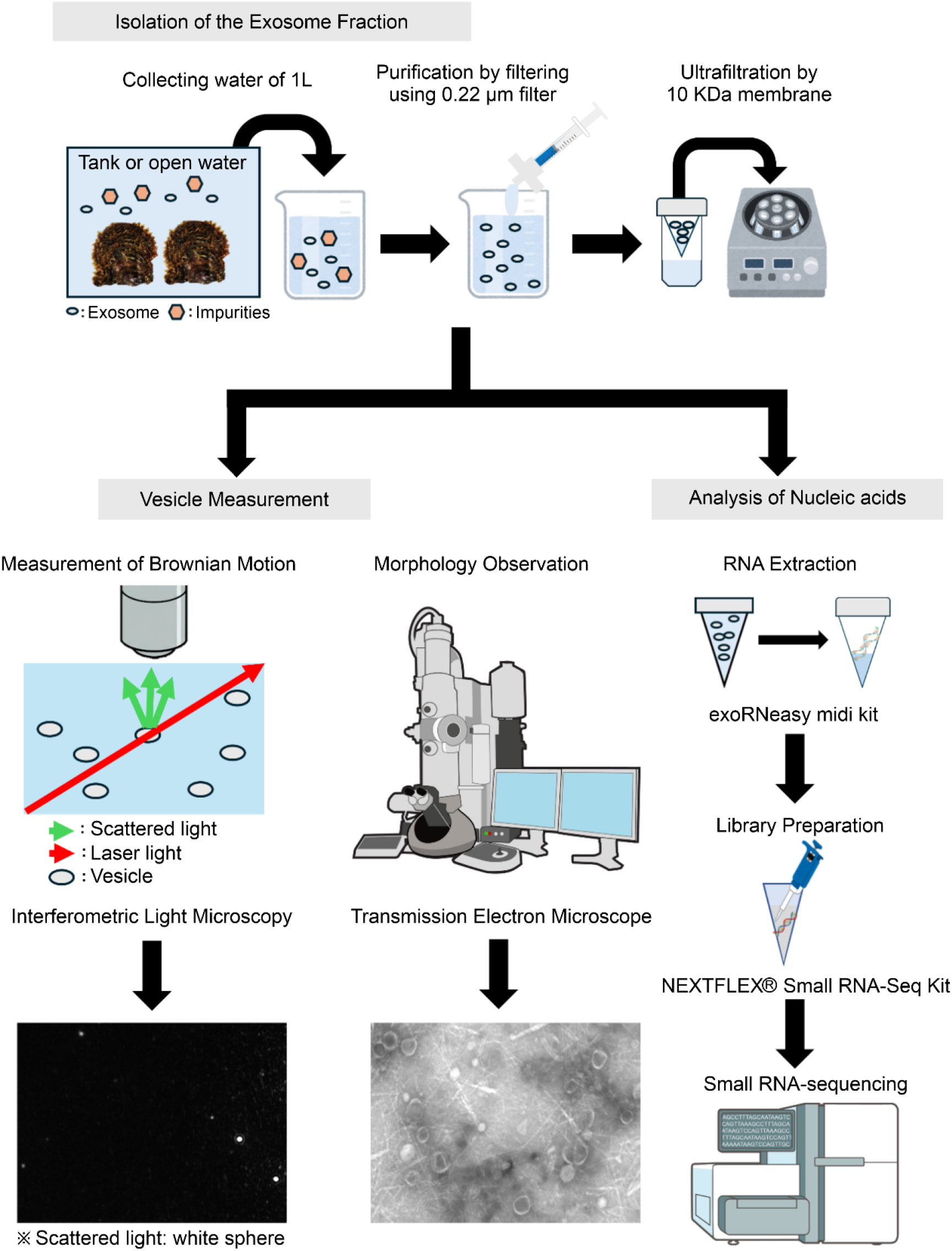
Experimental design and analysis workflow performed in this study.

Ultrafiltration was performed on 1 L of sample using Centricon Plus-70 10 kDa Ultracel-PL units (Merck Millipore) at 3,500 × g for over 25 min (Centrifuge; Model 6200; rotor: SF-2504S; KUBOTA, Tokyo, Japan). After removing the flow-through, the retained fraction was replenished with the remaining sample and repeatedly concentrated. The final concentrate was replaced with phosphate buffered saline (PBS) and stored at −80 °C until further use. For the aquaculture open-water samples, vesicle concentration and particle size distribution were analyzed using interferometric light microscopy (Videodrop), which measures Brownian motion (*61*). For this analysis, 100 μL samples from the morning and evening collections were sent to Meiwafosis (Tokyo, Japan). Vesicle counts per liter were estimated based on these measurements.

For transmission electron microscopy (TEM), samples from both the rearing tank and aquaculture area were prepared. Exosome/sEV fractions were further purified via ultracentrifugation at 100,000 × g for 70 min at 4 °C, supernatant was removed, and the pellet was resuspended in PBS. Ultracentrifugation was performed using an Optima MAX-TL ultracentrifuge (Beckman Coulter, CA, USA) with a TLA-100.3 rotor (Beckman Coulter). Samples were applied to self-made carbon coated copper grids (200 mesh), allowed to adsorb for less than 5 min, and negatively stained with 2% uranyl acetate (Bio-Rad, CA, USA) for approximately 30 seconds. The grids were then visualized using a transmission electron microscope (JEM-1400plus; JEOL, Tokyo, Japan) operated at an accelerating voltage of 120 kV.

### RNA extraction, library construction, and small RNA sequencing

Following the methodology described by Pan et al. (*62*), small RNA was extracted from each exosome fraction obtained from tank water (250 μL) and open water (300 μL) utilizing the exoRNeasy midi kit (Qiagen), in accordance with the manufacturer’s protocol. The quantity and quality of the extracted small RNA were evaluated using the Qubit microRNA kit with the Qubit 2.0 fluorometer (Thermo Fisher Scientific, MA, USA) and the Agilent Small RNA kit with the Agilent 2100 Bioanalyzer (Agilent Technologies, CA, USA). Library construction was carried out using the NEXTFLEX® Small RNA-Seq Kit v4 (Revvity, MA, USA), following the manufacturer’s instructions. For the evening samples, the cDNA amplification step was modified to increase the number of PCR cycles to 32 in order to obtain sufficient cDNA yields. Final library quality was evaluated using the High-Sensitivity D1000 ScreenTape Kit on the Agilent 2200 TapeStation. High-quality libraries were sequenced using the DNBSEQ-G400 platform (BGI) with 100 bp paired-end reads by BGI JAPAN (Hyogo, Japan).

### Sequencing data analysis

Following the method described by Meng et al. (*32*), we first trimmed adapter sequences and removed low-quality reads. Reads outside the 18–40 nt range were filtered using TrimGalore (https://github.com/FelixKrueger/TrimGalore). The remaining reads were mapped to the *P. fucata* reference genome (*63*) using Bowtie (*64*) with zero mismatches (bowtie -f -v 0 -a –al) to analyze expression and distribution across the genome. To improve alignment efficiency, reference haplotypes, versions 4.1A (reference) and 4.1B (alternative), were used. To classify and annotate small RNAs, mapped reads were compared against *Mollusca* sequences in the miRBase v22.1 database (*65*), allowing for one mismatch, using Bowtie (bowtie -v 1 -a --best –strata). The reads were also annotated against the Rfam 14.10 database (*66*) to identify other small RNA types (e.g., rRNAs, tRNAs), using Bowtie with no mismatches (bowtie -v 0 -a --best –strata). GNU Awk v4.0.2 (https://www.gnu.org/software/gawk/) was used to extract the sequences.

Subsequently, a custom Perl script (casify_rna.pl; Supplementary data 4), based on reference (*67*), was used to classify RNA and output FASTA sequences (perl classify_rna.pl -fa <fasta> -rfam <rfam_file> -mirna <miRNA.bwt> -ncrna <ncRNA.bwt> -outpre <output_prefix>). During annotation, sequences of unexpected lengths were observed among the classified miRNAs. Canonical miRNA sequences (20–24 nt) were extracted using seqkit (*68*); all others were designated as “unknown”. As Rfam lacks sequences specific to *P. fucata*, previously published sequences (28S rRNA: AB214477, 18S rRNA: AB214462, 5.8S rRNA: AB205102, ITS1: AB214218, ITS2: AB214265, rRNA intergenic spacer: AB214291 and AB214307) were used to recover unannotated rRNAs from the unknown category via local BLAST (BLAST 2.16.0+). Tandem repeats were then removed from the unknown sequences using MISA (*69*) and TRF (*70*). Sequences 25–32 nt in length were extracted and designated as predicted piRNAs, following the criteria in Huang et al. (*17*). In model organisms, piRNAs frequently show a 1U bias, with uridine at the first position (*33*–*35*). Accordingly, we analyzed the nucleotide composition at the first base of the predicted piRNAs, evaluating both the diversity and frequency of occurrence. Finally, we used piRNAs derived from *P. fucata* hemolymph exosomes (*17*) as a reference database to identify perfectly matching sequences via BLASTn (option: -task blastn-short).

Additionally, for the Tank and aquaculture site (AM1) sample, which yielded the highest number of sequencing reads, adapter trimming was performed using fastp (*71*). The resulting paired-end reads were mapped to the *P. fucata* reference genome using BWA-MEM (*72*), and alignment files were processed with SAMtools (*73*) to generate high-quality mapped reads. To account for potential splicing, alignments were also performed using HISAT2 (*74*), followed by transcript assembly with StringTie (*75*). A merged GTF file, generated from both genome versions 4.1A and 4.1B, was used to detect transcripts from both haplotypes. CDS and peptide sequences were predicted from the assembled transcripts using TransDecoder (*76*). Functional annotation was conducted via BLASTx and BLASTp searches against the Swiss-Prot database (downloaded December, 2023), and comprehensive annotations were generated using Trinotate (*77*), which incorporated protein domain and functional annotation data.

## Acknowledgments

We thank TOAGOSEI CO., LTD. and Mikimoto Pharmaceutical CO., LTD. for financial support, and the members of K. MIKIMOTO & CO., LTD. for their experimental assistance. We also express our sincere gratitude to Professor Yoshio Yamauchi, Department of Applied Biological Chemistry, Graduate School of Agricultural and Life Sciences, The University of Tokyo, Tokyo, Japan, for his valuable advice on ultracentrifugation methods. The authors wish to acknowledge the use of AI tools in the preparation of this manuscript. ChatGPT, GPT-5 (OpenAI) and Gemini, 2.5 pro (Google) were utilized to improve English grammar and clarity. These tools also provided assistance in generating and refining scripts for data analysis. The authors take full responsibility for the accuracy and originality of the content.

## Funding

This study was supported in part by Grant-in-Aid for Challenging Research (Pioneering) from the Japan Society for the Promotion of Science (JSPS) (25K21737; SA, RY), Core Research for Evolutional Science and Technology (CREST) from Japan Science and Technology Agency (JST) (JPMJCR23J2; NY).

## Author contributions

Conceptualization: RY, SA

Methodology: RY, LM, NH

Investigation: RY, LM, NH, II, SK

Formal analysis: RY, LM

Software: RY, LM, KY

Resources: NH, SM, KY, SK, KN, SA

Validation: TK, NY, NBK, TY, SW, SA

Visualization: RY, LM, II, NY, KN

Supervision: NY, TK, KM, KN, TY, SW, SA

Writing—original draft: RY

Writing—review & editing:RY, SA

## Competing Interests

The authors declare that they have no competing interests

## Data availability statement

The data have been deposited with links to BioProject accession number PRJDB37606 in the DNA Data Bank of Japan (DDBJ) BioProject database.

## MTA statement

Not applicable

## Supplementary Materials

**Supplementary data S1. (separate file)**

Complete list of estimated piRNA sequences and their read counts detected in the rearing tank and aquaculture samples.

**Supplementary data S2. (separate file)**

Complete list of occurrence counts of piRNA sequences matching known sequences in rearing tank sample.

**Supplementary data S3. (separate file)**

Summary of transcript assembly and annotation results generated by the Hisat2–StringTie– Trinotate pipeline, including transcript IDs, sample origin (Rearing tank/aquaculture; AM1), and BLASTx/BLASTp annotation data.

**Supplementary data S4 (separate file)**

Custom Perl script (classify_rna.pl) used to classify RNA types and generate corresponding FASTA sequence files.

Note: This file is named “Supplementary_data_4.pl”; please rename it to “classify_rna.pl” before use.

## Notes

### Competing Interest Statement

The authors have declared no competing interest.

